# Mitragynine and 7-Hydroxymitragynine: Bidirectional Effects on Breathing in Rats

**DOI:** 10.1101/2025.05.16.654392

**Authors:** Julio D. Zuarth Gonzalez, Alexandria K. Ragsdale, Sushobhan Mukhopadhyay, Christopher R. McCurdy, Lance R. McMahon, Samuel Obeng, Jenny L. Wilkerson

## Abstract

The use of kratom as an alternative to conventional opioids has surged, driven largely by anecdotal reports of its efficacy for pain relief and opioid withdrawal management. The growing prevalence of kratom products enriched with 7-hydroxymitragynine (7-HMG), an active metabolite of mitragynine (MG), necessitates evaluating the respiratory effects of these alkaloids and determining if naloxone reverses their potential respiratory depressant effects. Respiratory parameters were measured in awake, freely moving female and male Sprague-Dawley rats using whole body plethysmography. To minimize handling-induced artifacts and ensure precise respiratory recordings, drugs were administered intravenously. Morphine and 7-HMG induced significant respiratory depression, evidenced by reductions in breathing frequency, tidal volume, and minute volume. In contrast, MG administration unexpectedly increased respiratory frequency. Naloxone fully reversed the respiratory depression induced by both morphine and 7-HMG but did not alter the respiratory stimulant effects produced by MG. These findings demonstrate that 7-HMG exhibits significant respiratory depressant properties similar to classical opioids, and importantly, such depressant effects are effectively antagonized by naloxone. Conversely, MG exerts respiratory stimulant effects through mechanisms independent of opioid receptor pathways. Collectively, these data highlight crucial pharmacological distinctions between kratom alkaloids, underscoring the risk associated with high 7-HMG-containing kratom products and suggesting that the predominant alkaloid MG may offer a safer respiratory profile.

**Significance Statement:** The prevalence of kratom products containing 7-hydroxymitragynine (7-HMG), a µ-opioid receptor agonist, underscores the need to evaluate respiratory effects of kratom-related alkaloids and their reversal by naloxone. 7-HMG induced significant respiratory depression comparable to morphine, which was reversed by naloxone. Conversely, mitragynine, kratom’s most abundant alkaloid, unexpectedly increased respiratory frequency unaffected by naloxone. These findings highlight critical pharmacological differences between kratom-related alkaloids, emphasizing potential risks associated with products containing high concentrations of 7-HMG.

## Introduction

Opioids have long been recognized for their analgesic properties, yet their clinical utility is hindered by their potentially life-threatening side effects, like respiratory depression (Pattinson, 2008; Dahan et al., 2010; Boom et al., 2012). Opioid-induced respiratory depression (OIRD) represents a significant risk in both clinical settings and abuse-related overdoses (Volkow and McLellan, 2016; Overbeek et al., 2019). Preclinical studies have elucidated the mechanisms underlying OIRD, implicating μ-opioid receptor (MOR) activation in brainstem respiratory centers (Montandon and Slutsky, 2019; Varga et al., 2020). Clinical research has further highlighted the risk of OIRD in vulnerable populations such as the elderly and those with sleep-disordered breathing (MacIntyre et al., 2011; Gupta et al., 2018).

These safety concerns have driven the search for alternative analgesics with improved respiratory profiles. Kratom (*Mitragyna speciosa*) has garnered increasing attention for its opioid-like effects and potential therapeutic applications. In recent years, the use of kratom as an alternative to conventional opioids within the public domain has surged, driven by anecdotal reports of efficacy in managing pain and mitigating opioid withdrawal symptoms (Swogger and Walsh, 2018; Coe et al., 2019; Grundmann et al., 2024). Beyond these anecdotal reports, scientific interest in kratom continues to grow, with several studies focusing on its primary alkaloid, mitragynine (MG), and its hydroxylated metabolite, 7-hydroxymitragynine (7-HMG).

Although 7-HMG and MG were both shown to function as biased agonists at the MOR (Kruegel et al., 2016; Varadi et al., 2016; Faouzi et al., 2020; Gutridge et al., 2020), this does not mean that both 7-HMG and MG have fewer side effects than conventional opioids. Compounds developed as biased MOR agonists have failed to demonstrate improved safety profiles (Hill et al., 2018; Negus and Freeman, 2018; Kliewer et al., 2019; Kliewer et al., 2020).

Pharmacological characterization of the abuse liability of MG and 7-HMG revealed striking differences. MG did not substitute for morphine in the drug discrimination assay and was not self-administered, whereas 7-HMG fully substituted for morphine and demonstrated reinforcing effects (Harun et al., 2015; Yue et al., 2018; Hemby et al., 2019; Obeng et al., 2021). Additionally, 7-HMG *in vivo* has been demonstrated to be 4.4-to 13-fold more potent than morphine (Takayama et al., 2002; Matsumoto et al., 2004; Matsumoto et al., 2006; Kruegel et al., 2016; Obeng et al., 2021). These findings suggest that 7-HMG, but not MG, has greater liability risk and abuse potential similar to other opioid agonists despite biased agonism.

The differential abuse potential between these alkaloids is particularly concerning given the current market trends. A significant concern is the increasing availability of kratom products that contain high concentrations of 7-HMG (Lydecker et al., 2016; Smith et al., 2025). While analyses indicate that 7-HMG is either not detectable or present in extremely small amounts (<0.1%) in the living plant (Kikura-Hanajiri et al., 2009; Todd et al., 2020), likely formed as a post-harvest artifact (Chear et al., 2021), some manufacturers are marketing products containing 14–25 mg of 7-HMG per recommended serving, with labeled purities as high as 98% (Smith et al., 2025). This poses a risk, particularly for kratom-naïve individuals who may unknowingly purchase these products under the assumption that they are typical kratom preparations. These concerns underscore the need to evaluate the safety of both 7-HMG and MG, particularly regarding their effects on respiration.

The current state of knowledge regarding the respiratory effects of kratom and its related alkaloids remains limited and somewhat conflicting. One study found that oral MG did not yield significant dose-related respiratory depressant effects in rats, even at high doses (Henningfield et al., 2022). Another study in mice reported respiratory depression but observed a ceiling effect in MG-induced respiratory depression, while 7-HMG showed dose-dependent effects (Hill et al., 2022). However, the naloxone-reversibility of these respiratory effects remains to be determined. This is crucial as it would inform clinicians whether naloxone is effective in reversing 7-HMG induced respiratory depression.

In this study, we investigated the respiratory effects of the prototypical opioid, morphine, and kratom-related alkaloids (MG and 7-HMG) using whole body plethysmography in rats in two distinct experimental phases. In the first phase, we evaluated the direct respiratory effects of morphine, MG, and 7-HMG. In the second phase, we examined the potential for naloxone as a rescue intervention following MG and 7-HMG administration. The order of both the test compounds and naloxone rescue was counterbalanced between subjects. This investigation of the respiratory effects and naloxone-reversibility of these compounds provides new insights into the role of opioid receptor mechanisms in kratom-induced respiratory depression.

## Material and methods

### Subjects

A total of 32 Sprague-Dawley rats (Charles River, Houston, TX, USA; n=16 female, n=16 male), age 12 weeks, were used in this study. The rats were purchased with pre-implanted indwelling jugular catheters. For the first experimental phase, 16 animals were divided into two groups (n=8, morphine group; n=8, MG/7-HMG group). All testing followed a repeated-measure, within-subjects design where each animal received multiple doses of their respective drug(s). One male rat from the kratom-related alkaloids group expired before the completion of the experiment, resulting in a final sample size of n=7 for this group. The remaining 16 animals were used in the second experimental phase, with eight animals assigned to the morphine-naloxone group and eight animals assigned to the kratom-related alkaloids-naloxone group. Animals were housed individually within a temperature- and humidity-controlled vivarium and kept on a 12:12 light-dark cycle (lights on at 0700). Food and water were provided ad libitum. This study was conducted in accordance with the Guide for the Care and Use of Laboratory Animals (NRC, 2011) and was approved by the Institutional Animal Care Use Committee at Texas Tech University Health Sciences Center.

### Drugs

Morphine sulfate and naloxone hydrochloride (Fagron, St. Paul, MN) were diluted in normal saline. Mitragynine (MG) HCl and 7-hydroxymitragynine (7-HMG) were isolated as previously described (Ponglux et al., 1994) and were diluted in a vehicle solution consisting of 2.5% Tween 20, 2.5% Tween 80, 25% Polyethylene glycol, 25% Propylene glycol, and 45% normal saline. A mixture of heparin (30 units/ml) and gentamicin (2 mg/kg), diluted in normal saline, was used for antibiotic and anticoagulant prophylaxis.

### Apparatus

A whole body plethysmography system (Data Sciences International, St. Paul, MN) was used to measure and record the respiratory parameters in awake, freely moving rats. The system consisted of eight cylindrical acrylic chambers measuring 18.4 cm in diameter and 13.1 cm in height. The top interior portion of the chamber houses an infusion tether that is connected magnetically to the implanted vascular access button on the rats. The exterior portion of the chamber is connected to a pinport which facilitates aseptic intravenous (i.v.) drug delivery (PNP3M, Instech laboratories, Plymouth, PA, USA). The i.v. route was chosen to eliminate handling-induced respiratory artifacts during drug delivery, ensuring undisturbed respiratory recordings while enabling maximal effects with lower drug doses. Similarly, white noise was continuously played in the testing room to mask external sounds and minimize respiratory disturbances that could be caused by environmental noises.

The whole body plethysmography system measures box flow, which refers to the airflow in and out of the chamber caused by the animal’s respiration. When an animal inhales, air flows from the chamber into the animal’s respiratory system, creating a negative pressure within the chamber, which registers as a negative box flow. Conversely, during exhalation, air flows from the animal’s respiratory system back into the chamber, creating a positive pressure within the chamber, which registers as a positive box flow. A pressure transducer detects changes in box flow in the chamber which is airtight to prevent leakage. The pressure transducer converts pressure fluctuations caused by the animal’s breathing into electrical signals. These signals are then processed and analyzed by the whole body plethysmography software to calculate various respiratory parameters.

Respiratory rate (F, breaths/min) is directly measured from fluctuations in the positive and negative box flow. Tidal volume (TV, mL/breath) and minute volume (MV, mL/min) are estimated from the box flow signal. Tidal volume is equal to the product of the volume measured from the box flow and a compensation factor, which accounts for the animal’s body temperature, humidity, and barometric pressure. Minute volume is equal to the product of the volume measured from the box flow, respiratory rate, and compensation factor. In addition to these parameters, the system also calculates apneic pause, an index of respiratory constriction that represents decreased respiratory drive.

### Study Design

The study was conducted in two distinct experimental phases. In the first phase, we evaluated the direct respiratory effects by administering either saline or morphine (10 and 32 mg/kg, i.v.) in one group (n=8). A separate group (n=7) received either vehicle or the kratom-related alkaloids MG (5.6 and 10 mg/kg, i.v.) and 7-HMG (1, 3.2, and 10 mg/kg, i.v.). The order of drug administration was counterbalanced between subjects in both groups. For MG, we observed stimulatory effects but no respiratory depression at doses up to 10 mg/kg. Based on these findings, all animals in the kratom alkaloid group received a higher 17.8 mg/kg (i.v.) MG dose to assess potential respiratory depression at higher doses. This highest dose still produced no significant respiratory depression and resulted in seizure-like activity in two out of three male subjects, with one animal sustaining physical injury. This adverse reaction established 10 mg/kg as our upper safety limit for subsequent testing.

In the second phase, using entirely different animals, we examined the extent to which the opioid antagonist naloxone could reverse respiratory effects. The doses used in this phase (32 mg/kg for morphine, 10 mg/kg for MG, and 10 mg/kg for 7-HMG) were selected based on their significant effects on respiratory parameters during the first phase. One cohort (n=8) underwent four treatment conditions in a fully counterbalanced Latin square design: saline followed by saline, saline followed by naloxone (1 mg/kg, i.v.), morphine (32 mg/kg, i.v.) followed by saline, and morphine (32 mg/kg, i.v.) followed by naloxone (1 mg/kg, i.v.). Another cohort (n=8) underwent six treatment conditions using Williams design: vehicle followed by saline, vehicle followed by naloxone (1 mg/kg, i.v.), MG (10 mg/kg, i.v.) followed by saline, MG (10 mg/kg, i.v.) followed by naloxone (1 mg/kg, i.v.), 7-HMG (10 mg/kg, i.v.) followed by saline, and 7-HMG (10 mg/kg, i.v.) followed by naloxone (1 mg/kg, i.v.). In all second-phase treatments, naloxone or saline was administered 10 min after the initial compound injection.

Dose selection for this study was determined through preliminary testing in our laboratory. For morphine, doses of 10 and 32 mg/kg (i.v.) were selected based on pilot studies demonstrating that higher doses were necessary to produce reliable respiratory depression under normal air conditions without CO_2_ challenge, which typically reveals respiratory depression at lower doses. For the kratom-related alkaloids, we implemented a systematic dose-escalation approach due to the absence of published intravenous administration data. For the naloxone antagonism studies, we selected a dose of 1 mg/kg (i.v.) to provide effective blockade of opioid receptor-mediated effects given the relatively high doses of the test compounds used in our paradigm.

Animals were given a one-week habituation to new housing conditions before experiments commenced. After habituation, rats were transported to an experimental room mimicking conditions of the housing room and were placed individually in a whole body plethysmography chamber for a 3-hr acclimation period. During this time, rats received two vehicle injections to familiarize them with the injection procedure before returning to the housing room. For experimental sessions, each weekly testing began with a 60-min acclimation period followed by a 20-min baseline recording. After delivery of test compounds, respiratory parameters were recorded continuously for 60 min in both phases. Injections were manually administered over 15-30 s, and infusion lines were flushed with saline to ensure complete drug delivery.

To maintain catheter patency and prevent infections or blood clots, catheters were flushed with a solution containing gentamicin (4 mg/ml) and heparin (30 IU per ml) following catheter manipulation and at least every three days (Xi et al., 2008). On the day after each test session (Fridays), catheter patency was verified by infusing 0.2 ml of a solution containing ketamine (15 mg/ml) and midazolam (0.5 mg/ml) into the catheter (Zlebnik et al., 2014). Loss of righting reflex within 5 s indicated a patent catheter.

### Statistical Analysis

The study timeline consisted of 60 min of acclimation to whole body plethysmography chambers, followed by a 20-min baseline recording period. The acclimation period was neither plotted nor included in any analyses. After baseline recording, test compounds were administered intravenously, and post-drug measurements continued for 60 min, with data collected in 5-min bins. To account for individual differences and associated baseline measures, all respiratory parameters (frequency, tidal volume, and minute volume) were normalized by expressing each post-drug value as a percentage of the corresponding pre-drug baseline value. This normalization is particularly important for tidal volume, which is directly influenced by mass across species (Kleinman and Radford, 1964; Leith, 1983; Gomes et al., 2000; Quindry et al., 2016). For each experimental session, baseline values were calculated as the average of the 20-min recording period before drug infusion. For quantitative analysis, the time course data were transformed to area under the curve (AUC) values using the trapezoidal method, calculated over two specific time intervals: 0-30 minutes and 0-60 minutes post-administration.

For single-drug experiments with morphine, MG, and 7-HMG, AUC values were analyzed using one-way repeated measures ANOVAs separately for each respiratory parameter at the 30-minute and 60-minute intervals. Greenhouse-Geisser correction was applied to adjust degrees of freedom when the assumption of sphericity was violated. Following significant main effects, Dunnett’s multiple comparisons test was employed to compare drug treatment groups against the vehicle control.

In the naloxone antagonism experiments, morphine, MG, or 7-HMG administration was followed by naloxone or saline 10 minutes later. AUC values were calculated as described above and analyzed using two-way repeated measures ANOVAs with naloxone treatment and drug treatment as factors. The analysis examined the main effects of naloxone treatment and drug treatment, and their interaction at both time intervals. Tukey’s multiple comparisons test was used for multiple comparisons between treatment conditions. Statistical significance was established at p < 0.05 for all analyses. All statistical procedures were performed using GraphPad Prism (version 10) software.

## Results

### Respiratory Effects of Morphine, Mitragynine, and 7-Hydroxymitragynine Alone

Morphine produced dose-dependent respiratory depression, as illustrated by time courses for respiratory frequency, tidal volume, and minute volume (**Figure 1A-C**). Morphine had no significant effect on respiratory frequency at either 30 min [**Figure 1D**; F(1.57, 11.0) = 3.65, *p* = 0.069] or 60 min [**Figure 1E**; F(1.58, 11.1) = 3.06, *p* = 0.096]. Morphine significantly affected tidal volume at both 30 min [**Figure 1F**; F(1.25, 8.76) = 10.9, *p* = 0.007] and 60 min [**Figure 1G**; F(1.35, 9.42) = 16.9, *p* = 0.001]. Post hoc analysis revealed that only the higher dose of morphine (32 mg/kg) significantly reduced tidal volume at both time points (*p* < 0.001). Similarly, morphine significantly decreased minute volume at both 30 min [**Figure 1H**; F(1.31, 9.15) = 8.79, *p* = 0.012] and 60 min [**Figure 1I**; F(1.21, 8.44) = 9.73, *p* = 0.011]. Post hoc analysis showed that only the higher dose of morphine (32 mg/kg) significantly reduced minute volume at both time points (*p* < 0.001), while the lower dose (10 mg/kg) did not produce significant respiratory depression.

**Figure 1:**
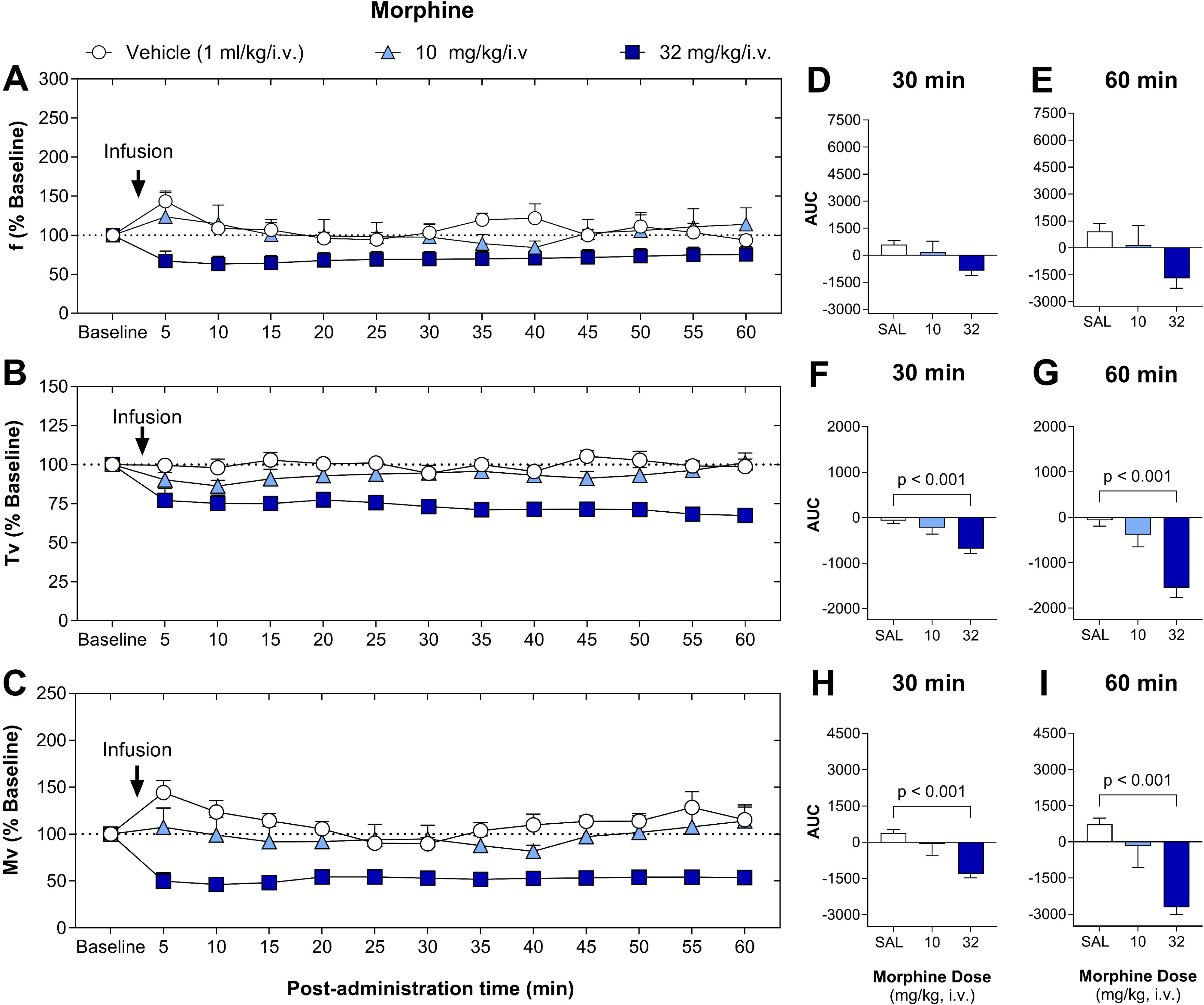
Time-course and AUC of morphine effects on respiratory parameters in rats. Respiratory frequency (*upper panels*), tidal volume (*middle panels*), and minute ventilation (*lower panels*) following intravenous administration of morphine in rats. Morphine was administered at doses of 10 mg/kg and 32 mg/kg, with saline (1 mL/kg) as the control. Data points for the time-course (A-C) represent the mean ± SEM (n=8), recorded in 5-minute bins continuously for up to 60 minutes post-infusion. Data for AUC are shown as mean ± SEM (n=8) for 30 (D, F, and H) and 60 (E, G, and I) minutes post-infusion. Statistically significant differences between saline and different doses are indicated (*p*-values).

MG administration produced distinct respiratory effects distinct from morphine, as shown in the time courses for frequency, tidal volume, and minute volume (**Figures 2A-C**). MG significantly increased respiratory frequency at both 30 min [**Figure 2D**; F(1.99, 11.9) = 4.00, *p* = 0.047] and 60 min [**Figure 2E**; F(1.78, 10.7) = 4.77, *p* = 0.036]. Post hoc analysis revealed that the 10 mg/kg dose significantly elevated respiratory frequency compared to vehicle control at both 30 min (*p* = 0.035) and 60 min (*p* = 0.046). MG did not significantly affect tidal volume at either 30 min [**Figure 2F**; F(1.53, 9.20) = 3.40, *p* = 0.086] or 60 min [**Figure 2G**; F(1.35, 8.09) = 3.93, *p* = 0.075]. Similarly, minute volume was not significantly altered by MG administration at either 30 min (**Figure 2H**; F(1.53, 9.17) = 4.07, *p* = 0.062) or 60 min (**Figure 2I**; F(1.59, 9.53) = 2.65, *p* = 0.128).

**Figure 2:**
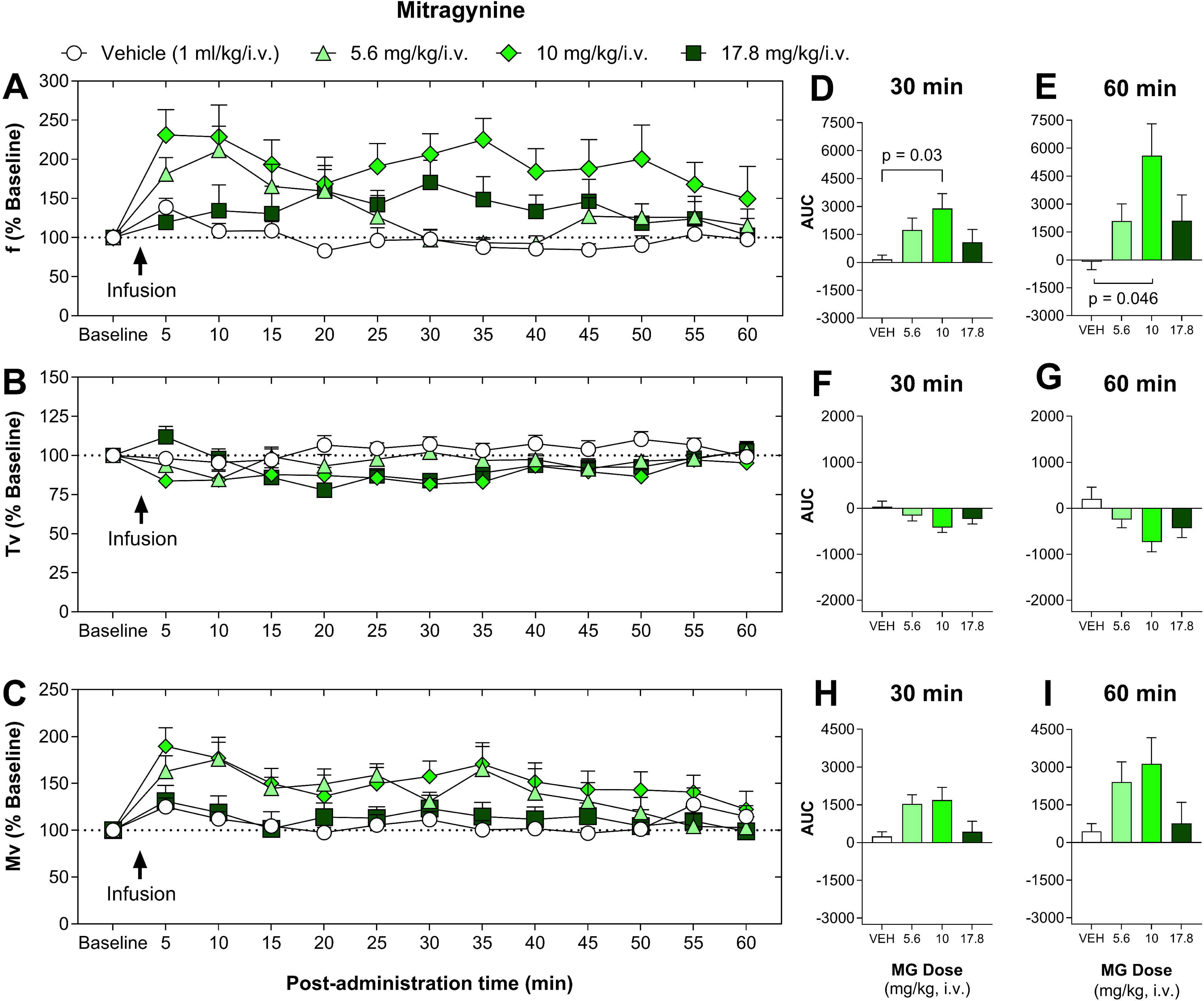
Time-course and AUC of MG effects on respiratory parameters in rats. Respiratory frequency (*upper panels*), tidal volume (*middle panels*), and minute ventilation (*lower panels*) following intravenous administration of MG in rats. MG was administered at doses of 5.6 mg/kg, 10 mg/kg, and 17.8 mg/kg, with vehicle (1 mL/kg) as the control. Data points for the time-course (A-C) represent the mean ± SEM (n=7), recorded in 5-minute bins continuously for up to 60 minutes post-infusion. Data for AUC are shown as mean ± SEM (n=8) for 30 (D, F, and H) and 60 (E, G, and I) minutes post-infusion. Statistically significant differences between vehicle and different doses are indicated (*p*-values).

Administration of 7-HMG produced robust, dose-dependent respiratory depression. Time courses for respiratory frequency, tidal volume, and minute volume are presented in **Figures 3A-C**. Respiratory frequency was significantly decreased by 7-HMG at both 30 min (**Figure 3D**; F(1.91, 11.5) = 12.7, *p* = 0.001) and 60 min (**Figure 3E**; F(2.26, 13.6) = 7.72, *p* = 0.005). Post hoc analysis revealed that both 3.2 mg/kg and 10 mg/kg doses significantly reduced respiratory frequency at 30 min (*p* = 0.04 and *p* = 0.03, respectively), but these effects were not maintained at 60 min. For tidal volume, 7-HMG approached but did not reach significance at both 30 min (**Figure 3F**; F(1.55, 9.31) = 4.19, *p* = 0.058) and 60 min (**Figure 3G**; F(1.57, 9.40) = 4.05, *p* = 0.061). Minute volume was significantly reduced by 7-HMG at both 30 min (**Figure 3H**; F(2.17, 13.0) = 17.9, *p* < 0.001) and 60 min (**Figure 3I**; F(2.35, 14.1) = 8.52, *p* = 0.003). Post hoc analysis revealed that the highest doses (3.2 and 10 mg/kg/i.v.) significantly decreased minute volume at 30 min (*p* = 0.005 for both doses), but these effects were not statistically significant at 60 min.

**Figure 3:**
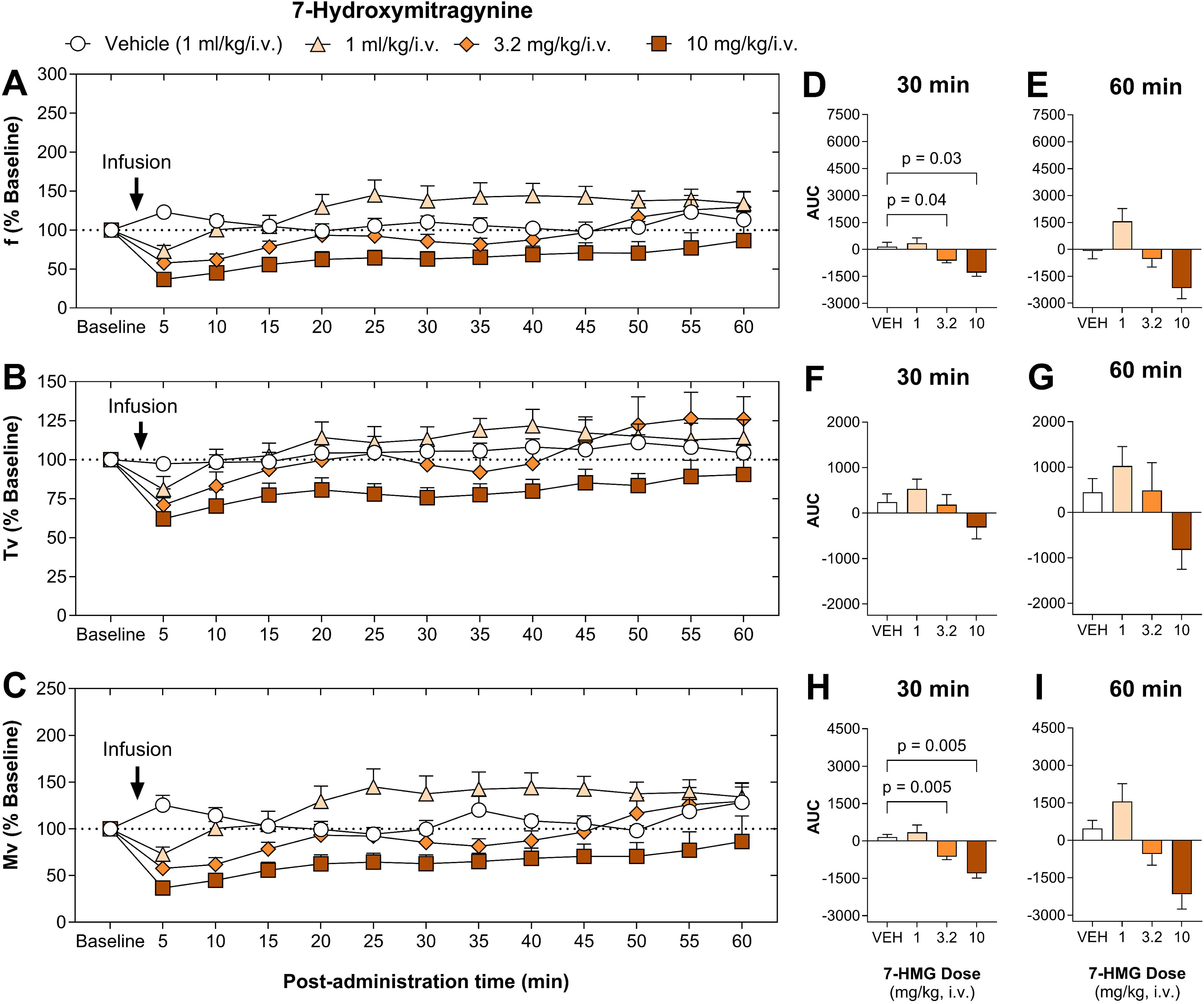
Time-course and AUC of 7-HMG effects on respiratory parameters in rats. Respiratory frequency (*upper panels*), tidal volume (*middle panels*), and minute ventilation (*lower panels*) following intravenous administration of 7-HMG in rats. 7-HMG was administered at doses of 1 mg/kg, 3.2 mg/kg, and 10 mg/kg, with vehicle (1 mL/kg) as the control. Data points for the time-course (A-C) represent the mean ± SEM (n=7), recorded in 5-minute bins continuously for up to 60 minutes post-infusion. Data for AUC are shown as mean ± SEM (n=8) for 30 (D, F, and H) and 60 (E, G, and I) minutes post-infusion. Statistically significant differences between vehicle and different doses are indicated (*p*-values).

## Naloxone Antagonism Studies

### Morphine + Naloxone

Naloxone (1 mg/kg) treatment effectively reversed the respiratory depression induced by morphine (32 mg/kg). Time courses for respiratory frequency, tidal volume, and minute volume are presented in **Figures 4A-C**. For respiratory frequency, two-way repeated measures ANOVA at 30 min revealed significant main effects of naloxone [F(1, 7) = 7.02, *p* = 0.033], morphine [F(1, 7) = 6.24, *p* = 0.041], and naloxone × morphine interaction [F(1, 7) = 21.9, *p* = 0.002] (**Figure 4D**). Despite a significant main effect of naloxone, there was no significant difference between saline + naloxone and saline + saline control. Additionally, multiple comparisons revealed that neither morphine + saline nor morphine + naloxone administration significantly impacted respiratory frequency compared to control, consistent with the results of the morphine dose-response experiment. However, the direct comparison between morphine + saline and morphine + naloxone treatments revealed a statistically significant difference (*p* = 0.02). At 60 min, these effects persisted with significant main effects of naloxone [F(1, 7) = 9.98, *p* = 0.016], morphine [F(1, 7) = 8.39, *p* = 0.023], and naloxone × morphine interaction [F(1, 7) = 23.8, *p* = 0.002] (**Figure 4E**). At this time point, morphine + naloxone was not only significantly different from morphine + saline (*p* = 0.02) but also significantly increased compared to saline control (*p* = 0.04), demonstrating an overshoot effect.

**Figure 4:**
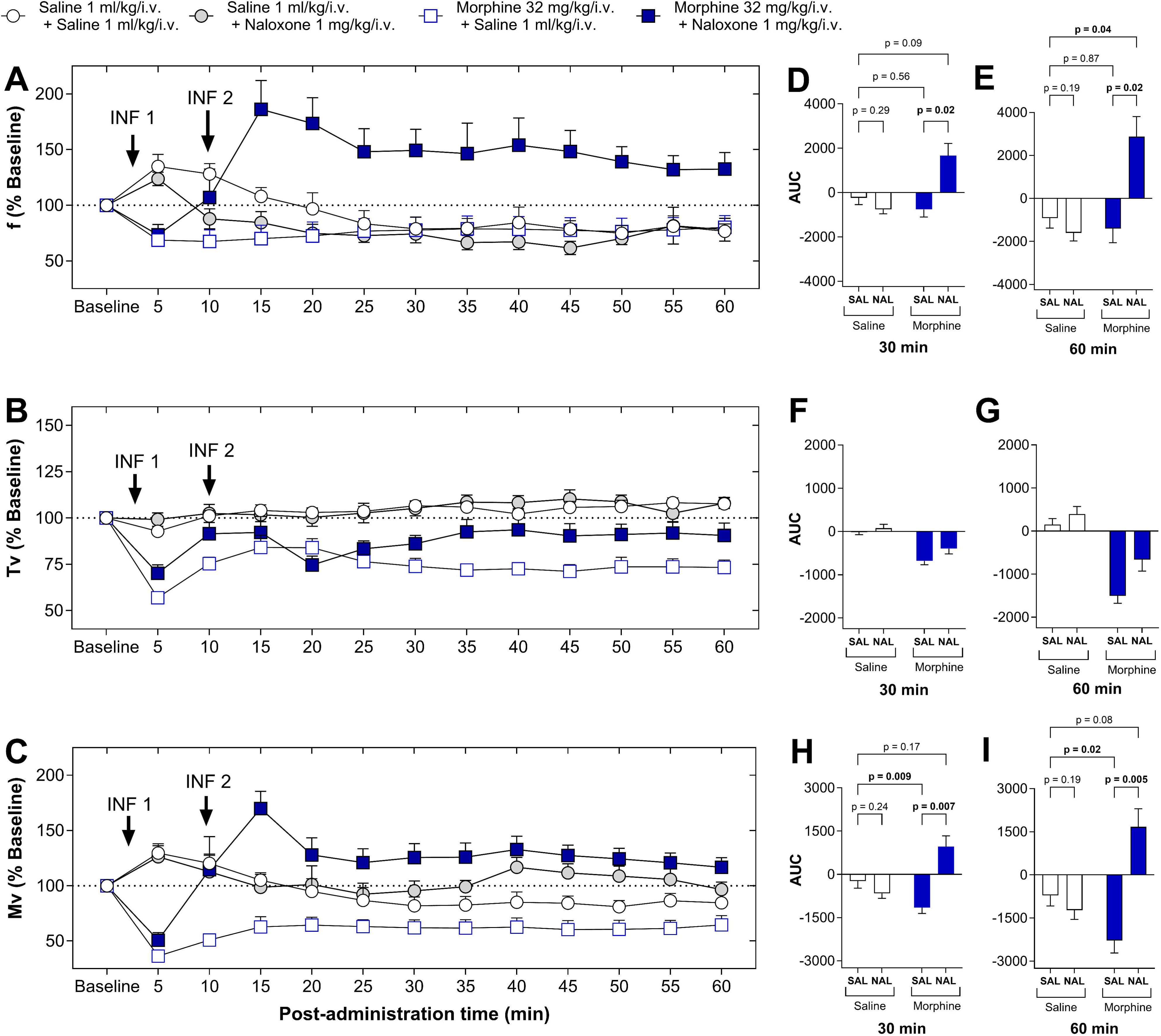
Time-course and AUC of morphine and naloxone effects on respiratory parameters in rats. Respiratory frequency (*upper panels*), tidal volume (*middle panels*), and minute ventilation (*lower panels*) following intravenous administration of morphine with naloxone rescue treatment in rats. Morphine was administered (32 mg/kg) followed by naloxone (1 mg/kg) rescue treatments after 10 minutes. Data points for the time-course (A-C) represent the mean ± SEM (n=7), recorded in 5-minute bins continuously for up to 60 minutes post-morphine infusion. Data for AUC are shown as mean ± SEM (n=8) for 30 (D, F, and H) and 60 (E, G, and I) minutes post-morphine infusion. Statistically significant differences between treatments are indicated (*p*-values).

For tidal volume at 30 min, a significant effect of morphine [F(1, 7) = 27.5, *p* = 0.001] was observed, while naloxone [F(1, 7) = 3.57, *p* = 0.101] and naloxone × morphine interaction [F(1, 7) = 1.54, *p* = 0.254] did not reach significance (**Figure 4F**). At 60 min, both naloxone [F(1, 7) = 7.22, *p* = 0.031] and morphine [F(1, 7) = 27.8, *p* = 0.001] showed significant effects but not the naloxone x morphine interaction (F(1, 7) = 3.28, *p* = 0.113] (**Figure 4G**). No post-hoc analyses were conducted for tidal volume due to the absence of significant interactions at either time point.

For minute volume at 30 min, two-way repeated measures ANOVA showed significant effects of naloxone [F(1, 7) = 9.76, *p* = 0.017] and naloxone × morphine interaction [F(1, 7) = 41.0, *p* < 0.001] (**Figure 4H**). Post-hoc analysis revealed significant depression with morphine alone compared to control (*p* = 0.009), which was not significant when naloxone was co-administered. Additionally, morphine + saline and morphine + naloxone treatments were significantly different from each other (*p* = 0.007). At 60 min, significant effects of naloxone [F(1, 7) = 15.5, *p* = 0.006] and naloxone × morphine interaction [F(1, 7) = 45.0, *p* < 0.001] persisted (**Figure 4I**). The same pattern of post-hoc differences was observed at this timepoint, with morphine alone causing significant depression compared to control (*p* = 0.02), while morphine + naloxone was not significantly different from control but was significantly different from the morphine + saline group (*p* = 0.005).

### Mitragynine + Naloxone

Administration of naloxone failed to antagonize MG’s respiratory effects. Time courses for respiratory frequency, tidal volume, and minute volume are presented in **Figure 5A-C**, respectively. Two-way repeated measures ANOVA for respiratory frequency showed that MG had a significant effect at both 30 min [**Figure 5D**; F(1, 7) = 14.4, *p* = 0.007] and 60 min [**Figure 5E**; F(1, 7) = 8.99, *p* = 0.020], while naloxone effects and interactions were not significant at either timepoint. Despite the lack of naloxone effects, the significant main effect of MG indicates that both MG + saline and MG + naloxone treatments resulted in elevated respiratory frequencies compared to controls.

**Figure 5:**
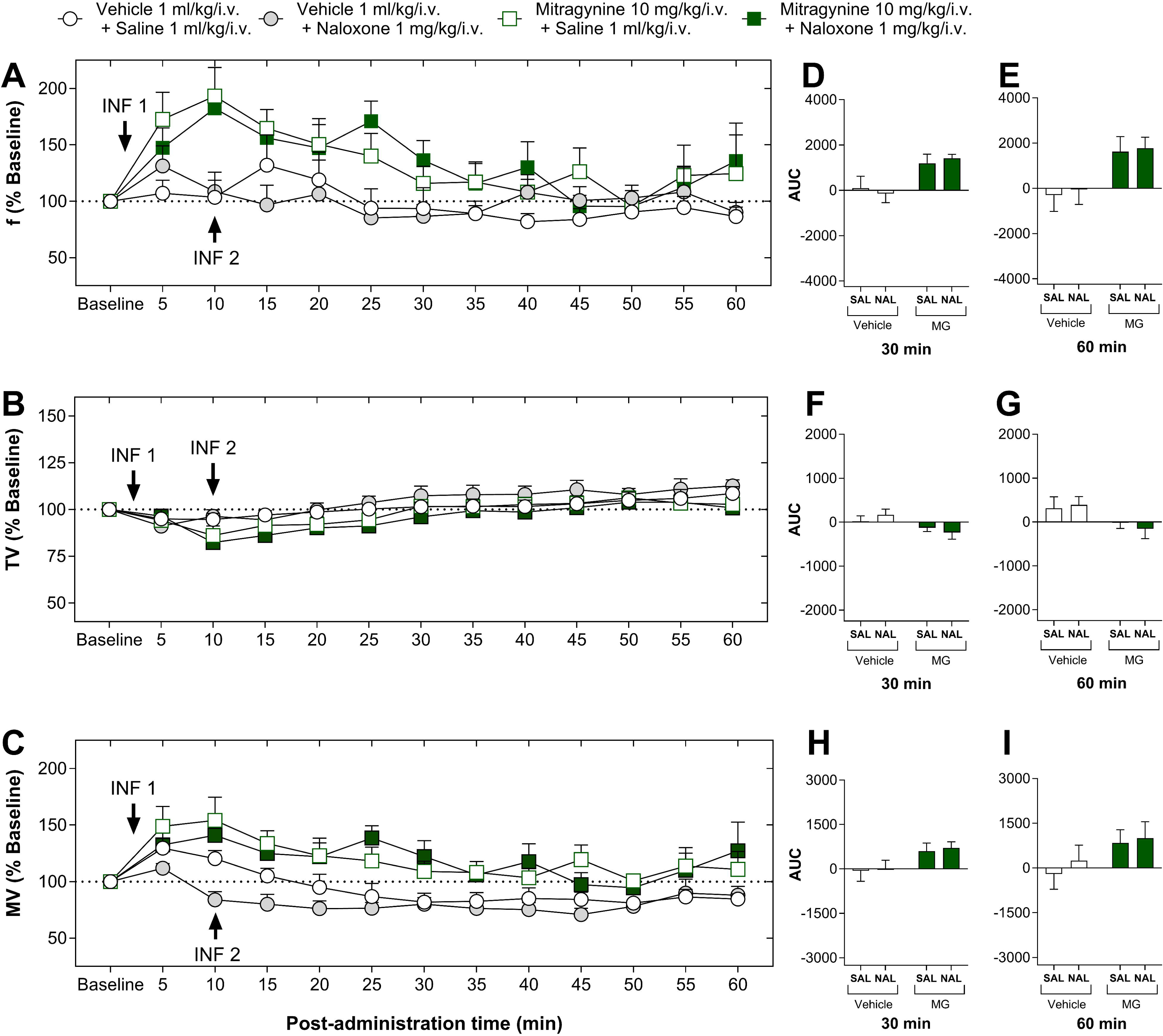
Time-course and AUC of MG and naloxone effects on respiratory parameters in rats. Respiratory frequency (*upper panels*), tidal volume (*middle panels*), and minute ventilation (*lower panels*) following intravenous administration of MG with naloxone rescue treatment in rats. MG was administered (10 mg/kg) followed by naloxone (1 mg/kg) rescue treatments after 10 minutes. Data points for the time-course (A-C) represent the mean ± SEM (n=8), recorded in 5-minute bins continuously for up to 60 minutes post-MG infusion. Data for AUC are shown as mean ± SEM (n=8) for 30 (D, F, and H) and 60 (E, G, and I) minutes post-MG infusion. Statistically significant differences between treatments are indicated (*p*-values).

For tidal volume, MG showed a near-significant effect at 30 min [**Figure 5F**; F(1, 7) = 5.26, *p* = 0.055] and a significant effect at 60 min [**Figure 5G**; F(1, 7) = 7.18, *p* = 0.032], with no significant naloxone effects or interactions. For minute volume, two-way repeated measures ANOVA showed no significant effects of MG, naloxone, or their interaction at either 30 min (**Figure 5H**) or 60 min (**Figure 5I**). These results indicate that naloxone did not effectively antagonize the respiratory effects of MG, suggesting a mechanism distinct from typical opioid activity.

### 7-Hydroxymitragynine + Naloxone

Naloxone effectively reversed the respiratory depression induced by 7-HMG. Time courses for respiratory frequency, tidal volume, and minute volume are presented in **Figures 6A**, **6B**, and **6C**, respectively. Two-way RM ANOVA for respiratory frequency at 30 min showed that while main effects of naloxone [F(1, 7) = 0.460, *p* = 0.519] and 7-HMG [F(1, 7) = 0.0922, *p* = 0.770] were not significant, a significant naloxone × 7-HMG interaction [F(1, 7) = 12.5, *p* = 0.010] was observed (**Figure 6D**). Further analysis of the interaction using multiple comparisons demonstrated that neither 7-HMG + saline nor 7-HMG + naloxone administration significantly impacted respiratory frequency compared to control. The direct comparison between 7-HMG + saline and 7-HMG + naloxone further revealed a significant difference (*p* = 0.03). At 60 min, no significant main effects or interactions were detected (**Figure 6E**).

**Figure 6:**
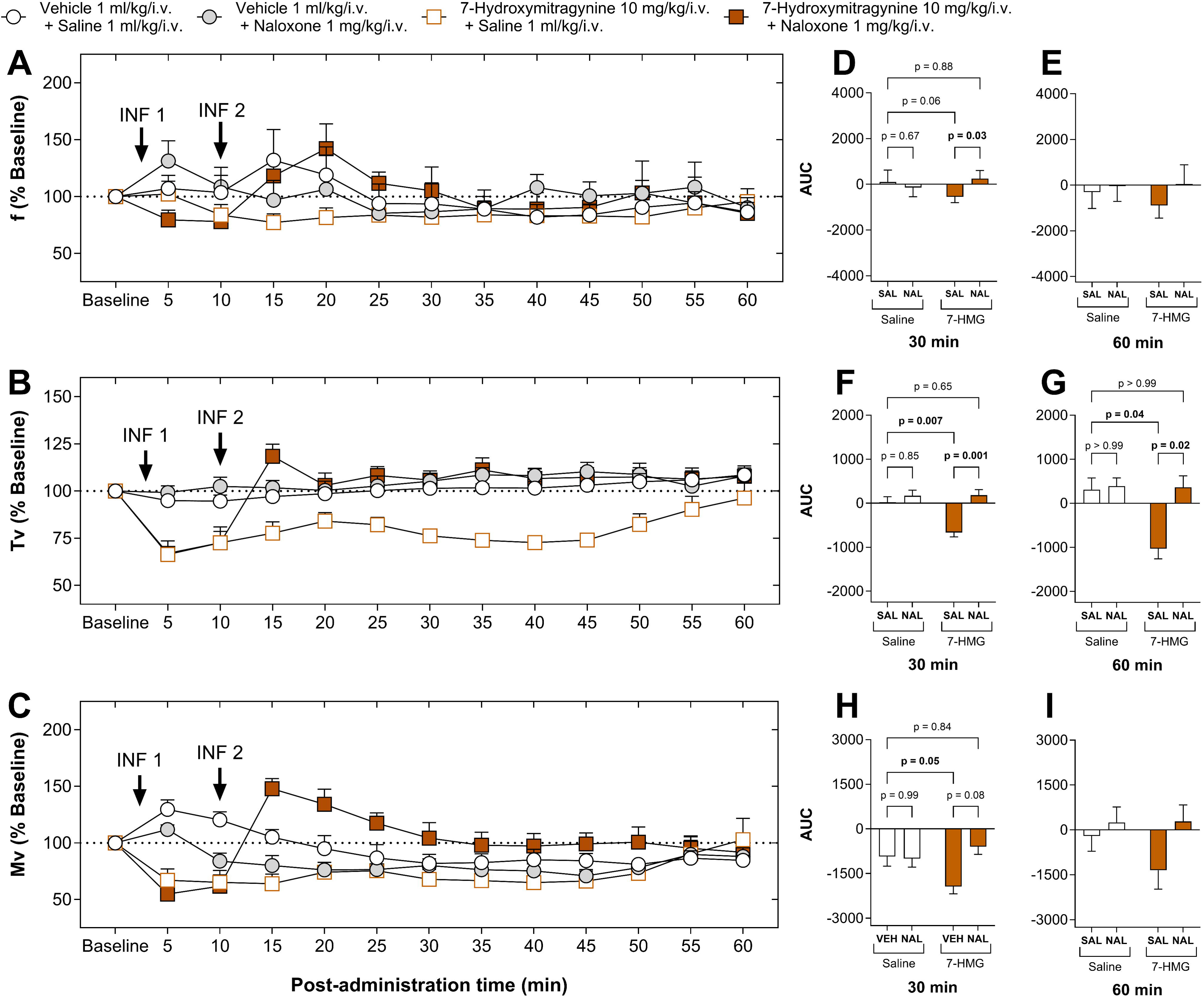
Time-course and AUC of 7-HMG and naloxone effects on respiratory parameters in rats. Respiratory frequency (*upper panels*), tidal volume (*middle panels*), and minute ventilation (*lower panels*) following intravenous administration of 7-HMG with naloxone rescue treatment in rats. 7-HMG was administered (10 mg/kg) followed by naloxone (1 mg/kg) rescue treatments after 10 minutes. Data points for the time-course (A-C) represent the mean ± SEM (n=8), recorded in 5-minute bins continuously for up to 60 minutes post-7-HMG infusion. Data for AUC are shown as mean ± SEM (n=8) for 30 (D, F, and H) and 60 (E, G, and I) minutes post-7-HMG infusion. Statistically significant differences between treatments are indicated (*p*-values).

For tidal volume, two-way RM ANOVA demonstrated significant effects of naloxone [F(1, 7) = 18.0, *p* = 0.004], 7-HMG [F(1, 7) = 10.4, *p* = 0.014], and naloxone × 7-HMG interaction [F(1, 7) = 11.7, *p* = 0.011] at 30 min (**Figure 6F**). Despite a significant main effect of naloxone, there was no significant difference between vehicle + naloxone and saline + saline control. Post-hoc analysis also revealed significant depression with 7-HMG + saline compared to control (*p* = 0.007), which was not observed when naloxone was co-administered (*p* = 0.65). Additionally, the 7-HMG + saline and 7-HMG + naloxone treatments were significantly different from each other (*p* = 0.001). These effects on tidal volume persisted at 60 min for naloxone [F(1, 7) = 18.0, *p* = 0.004], 7-HMG [F(1, 7) = 10.4, *p* = 0.014] and naloxone × 7-HMG interaction [F(1, 7) = 11.7, *p* = 0.011] (**Figure 6G**). At 60 min, 7-HMG + saline showed significant depression compared to control (*p* = 0.04), while the 7-HMG + naloxone group was not significantly different from control. The difference between 7-HMG + saline and 7-HMG + naloxone treatments remained significant (*p* = 0.02).

For minute volume, despite non-significant main effects of naloxone [F(1, 7) = 3.85, *p* = 0.090] and 7-HMG [F(1, 7) = 1.58, *p* = 0.249] at 30 min, a significant naloxone × 7-HMG interaction [F(1, 7) = 26.7, *p* = 0.001] was observed (**Figure 6H**). Post-hoc analysis at 30 min revealed a significant minute volume depression in the 7-HMG + saline group compared to control (*p* = 0.05), which was non-significant when naloxone was also administered. At 60 min, the interaction approached but did not reach significance [**Figure 6I**; F(1, 7) = 5.07, *p* = 0.059].

## Discussion

In the present study, both morphine and 7-HMG produced significant respiratory depression as indicated by decreases in breathing frequency, tidal volume and minute volume, while MG unexpectedly increased respiratory frequency. Naloxone significantly reversed (i.e., antagonized) the respiratory depressant effects of both morphine and 7-HMG, while the respiratory stimulant effects of MG were not antagonized or otherwise modified by naloxone administration. These results provide strong evidence that 7-HMG shares respiratory depressant effects with classical opioids, whereas MG increases respiration through mechanisms not involving opioid receptors.

Morphine, our reference compound, exhibited clear respiratory depression evidenced by decreases in tidal volume and minute volume. At the highest dose tested (32 mg/kg), morphine significantly reduced tidal volume and minute volume, while no significant effects were observed at the lower dose (10 mg/kg), indicating dose-dependent respiratory depression under ambient air conditions. These effects were completely reversed by naloxone, confirming the well-established role of opioid receptor mechanisms in morphine-induced respiratory depression (Longnecker et al., 1973; McGilliard and Takemori, 1978; Gairola et al., 1980). Our findings are consistent with previously documented morphine-induced decreases in respiration across multiple species using various routes of administration (Flórez et al., 1972; Thompson et al., 1995; Borgbjerg et al., 1996; Boom et al., 2012).

In the current study, MG did not produce respiratory effects typical of most opioid receptor agonists, i.e., did not decrease respiration up to 17.8 mg/kg. Doses of MG larger than 17.8 mg/kg produced overt signs of toxicity and were not fully tested. Instead, MG produced significant increases in respiratory frequency. In previously published studies, MG was reported to decrease some respiratory parameters (Hill et al., 2022). Lyophilized kratom tea also decreased respiration rates (Wilson et al., 2020). However, the experimental parameters of the previous studies (e.g., oral administration in mice) differed from the experimental parameters of the current study (e.g., intravenous administration in rats). With respect to route of administration, CYP450-mediated conversion of MG into its active metabolites is expected to be greater following oral as compared with intravenous administration (Hanapi et al., 2013; Basiliere and Kerrigan, 2020; Kamble et al., 2020; Kamble et al., 2023). While metabolism to sufficiently large amounts of 7-HMG could potentially contribute to the effects produced by MG administration, our group has demonstrated MG-induced antinociception in mice is evident when the amounts of 7-HMG measured in the brain are far less than the amounts required for antinociceptive effects (Berthold et al., 2022). Therefore, experimental support for a MG prodrug hypothesis is currently lacking, and any differences in 7-HMG formation across routes of administration do not appear to explain variable outcomes in studies of antinociception. Alternatively, the effects of MG appear to differ consistently between mice (Wilson et al., 2020; Hill et al., 2022) and rats (Henningfield et al., 2022). For example, under experimental conditions in rats that are highly sensitive to the antinociceptive effects of 7-HMG and other opioid agonists, our group has demonstrated that MG administration is devoid of antinociceptive effects (Obeng et al., 2021). The present study extends these findings to include respiratory effects, which in rats are not merely lacking but rather opposite in direction to the respiratory depressant effects of opioids.

Results for 7-HMG demonstrated significant dose-dependent respiratory depression, particularly at the 10 mg/kg dose. Importantly, we observed distinct time-dependent effects in the respiratory parameters. At 30 minutes post-administration, 7-HMG significantly decreased respiratory frequency and minute volume at both 3.2 mg/kg and 10 mg/kg doses, but these effects were not maintained at 60 minutes. This temporal profile indicates that 7-HMG’s respiratory depressant effects peak earlier and begin to diminish over time, contrasting with morphine which maintained effects across both 30 and 60 min. The reductions in respiratory frequency, tidal volume, and minute volume by 7-HMG observed in this study parallel the characteristic respiratory profile of classical μ-opioid agonists like morphine which are consistent with prior findings where 7-HMG showed respiratory depressant effects in mice (Hill et al., 2022). Notably, the reversal of 7-HMG-induced respiratory depression by naloxone provides pharmacological evidence for opioid receptor-mediated effects. The efficacy of naloxone antagonism, particularly evident in tidal volume measurements, supports substantial MOR involvement in 7-HMG’s respiratory effects. These findings align with previous *in vitro* characterization, demonstrating that 7-HMG has comparable binding affinity (K*i*) to morphine at μ-opioid receptors but 16-fold higher potency while only reaching less than half of morphine’s maximal efficacy (Emax) (Matsumoto et al., 2006; Kruegel et al., 2019; Obeng et al., 2020; Obeng et al., 2021). Collectively, these results suggest that 7-HMG functions primarily as a MOR agonist.

The opposing respiratory effects of MG and 7-HMG suggest that MG—the most abundant kratom alkaloid, constituting up to 66% of the total alkaloidal fraction (Raffa, 2014)— may counteract the respiratory depressive effects of 7-HMG. In the study by Hill et al., MG was shown to reduce minute volume; however, its respiratory depressant effects exhibited a ceiling effect, such that respiratory depression did not continue to worsen at higher doses of MG. In contrast, 7-HMG did not display a ceiling effect (Hill et al., 2022). Furthermore, Wilson et al. demonstrated that while lyophilized kratom tea (containing all the alkaloids in the plant) decreased respiratory rate, the reduction in respiration caused by morphine was significantly greater than that produced by the kratom tea (Wilson et al., 2020). These findings suggest that MG could mitigate the respiratory depressant effects of 7-HMG, given that the magnitude of 7-HMG-induced respiratory depression is similar to that of morphine. Taken together, these results support the idea that MG contributes to the improved safety profile of kratom. However, caution is warranted with higher doses of MG, as administration of a high dose of MG (17.8 mg/kg) induced seizure-like activity without causing respiratory depression. This observation aligns with poison control center data (2011-2017) documenting 113 seizures among 1174 kratom exposures (Fletcher and Dhyani, 2025).

Adding to these concerns is the rapid proliferation of kratom-associated products specifically marketed for their high 7-HMG content (Lydecker et al., 2016; Smith et al., 2025). Our examination of two major online retailers catalogs reveal a disturbing trend. At the time of submission, over 401 products specifically marketed for their 7-HMG content are readily available to consumers (Supplemental Tables 1-2). Although the range spans from 3.5 mg to an alarming 50 mg per serving, most products contain between 7-15 mg of 7-HMG per recommended dose, still considerably higher than levels found in traditional kratom preparations (Supplemental Table 1). Many products explicitly highlight their “ultra potent” formulations. These products are marketed in various consumer-friendly formats including tablets/pills, liquid shots/syrups, vape pens, sublingual strips, edible products, and even gummies, potentially appealing to kratom-naïve individuals (Supplemental Tables 1-2). Such formulations may be mistakenly perceived as typical kratom products, increasing the risk of unintentional exposure to pharmacologically significant doses of 7-HMG. The ability of 7-HMG to induce respiratory depression similar to morphine, highlights the urgent need for the regulation of these 7-HMG containing products.

While the findings of this study contribute significantly to our understanding of the respiratory effects of MG and 7-HMG and their interaction with naloxone, several limitations should be considered. First, we examined respiration after acute administration only; repeated administration might yield different respiratory profiles. Second, while we focused on respiratory parameters, a comprehensive safety evaluation would require assessment of additional physiological systems, such as cardiovascular and carcinogenic toxicity. Third, kratom contains numerous alkaloids; additional research is needed to determine the respiratory effects of other constituent alkaloids and their combinations. Future studies are also needed to investigate the molecular mechanisms underlying the distinct respiratory profile of MG, which was not affected by naloxone.

## Conclusions

These respiratory depression findings have direct implications for evaluating kratom’s safety profile and informing regulatory decisions. The observation that MG does not produce respiratory depression, and may even stimulate respiration, is particularly intriguing. However, the potent respiratory depressant effects of 7-HMG raise significant safety concerns, especially considering the increasing presence of novel semi-synthetic products that may contain up to 98% 7-HMG (Smith et al., 2025). Importantly, the respiratory depression produced by 7-HMG is sensitive to naloxone, which may have lifesaving implications in emergency medicine.

## Acknowledgments

This work was supported by National Institutes of Health National Institute on Drug Abuse [Grants DA48353 and UG3/UH3 DA048353 01]; Texas Tech University Health Sciences Center Office of Research and the Jerry H. Hodge School of Pharmacy. The authors would like to thank Dr. Renata Marchette and Dr. George Koob for providing training to JLW on the use of rat whole body plethysmography. The authors also gratefully acknowledge Dr. Aiden Hampson for his valuable input during the development of this project.

## Author Contributions

Conceptualization: JDZG, CRM, LRM, and JLW; Funding acquisition: CRM and LRM; Formal analysis: JDZG, SO, and JLW; Investigation: JDZG and AKR; Resources: SM and CRM; Supervision and Project administration: CRM, LRM, and JLW; Writing - Original draft: JDZG, SO, CRM, LRM, and JLW; Writing - Review & Editing; JDZG, AKR, SM, SO, CRM, LRM, and JLW. All authors have read, edited, and approved the final version of the manuscript.

## Data Availability Statement

The data that support the findings of this study are available within the paper. Any additional information is available on request from the corresponding authors.

## Nonstandard Abbreviations

AUC: Area under the curve
7-HMG: 7-Hydroxymitragynine
i.v.: Intravenous
MV: Minute volume
MG: Mitragynine
MOR: μ-opioid receptor
OIRD: Opioid-induced respiratory depression
F: Respiratory rate
TV: Tidal volume

